## Supplemental Material for "Benchmark dataset for training machine learning models to predict the pathway involvement of metabolites"

Table S1 – Parameters for configuring the atom coloring method.

| **Method Parameter** | **Parameter Description** | **Value We Chose** | **Reason For Value** |
|---|---|---|---|
| r_groups | If true, add R groups in the coloring. | True | Enabled us to replace 'R' symbols with 'C' (most R-groups are bonded to the rest of the molecule beginning with a carbon) and thus include more atom color detail. |
| bond_stereo | If true, add bond stereo detail when constructing colors. | True | Added stereochemistry detail to atom colors which is relevant to predicting pathway involvement and more precisely distinguishes compounds. |
| atom_stereo | If true, add atom stereo detail when constructing colors. | True | Added stereochemistry detail to atom colors which is relevant to predicting pathway involvement and more precisely distinguishes compounds. |
| resonance | If true, ignore the difference between double and single bonds. | False | Resonance set to True treats all bonds as single bonds whereas setting it to False distinguishes bond order. |
| isotope_resolved | If true, add isotope detail when constructing colors. | False | Isotope specification of atoms does not uniquely identify a compound since identical compounds can contain atoms with different isotopes. |

| | | | |
|---|---|---|---|
| charge | If true, add charge detail when constructing colors. | False | Atom charge does not uniquely identify a compound since the only difference is electron content rather than elemental identity. |
| backbone | If true, ignore bond types in the coloring. | False | Added bond-type detail to atom colors which is relevant to predicting pathway involvement and more precisely distinguishes compounds. |

Figure S1 – Distribution of mean minus median differences of feature importance scores for each pathway category in the full dataset trained on the XGBoost model

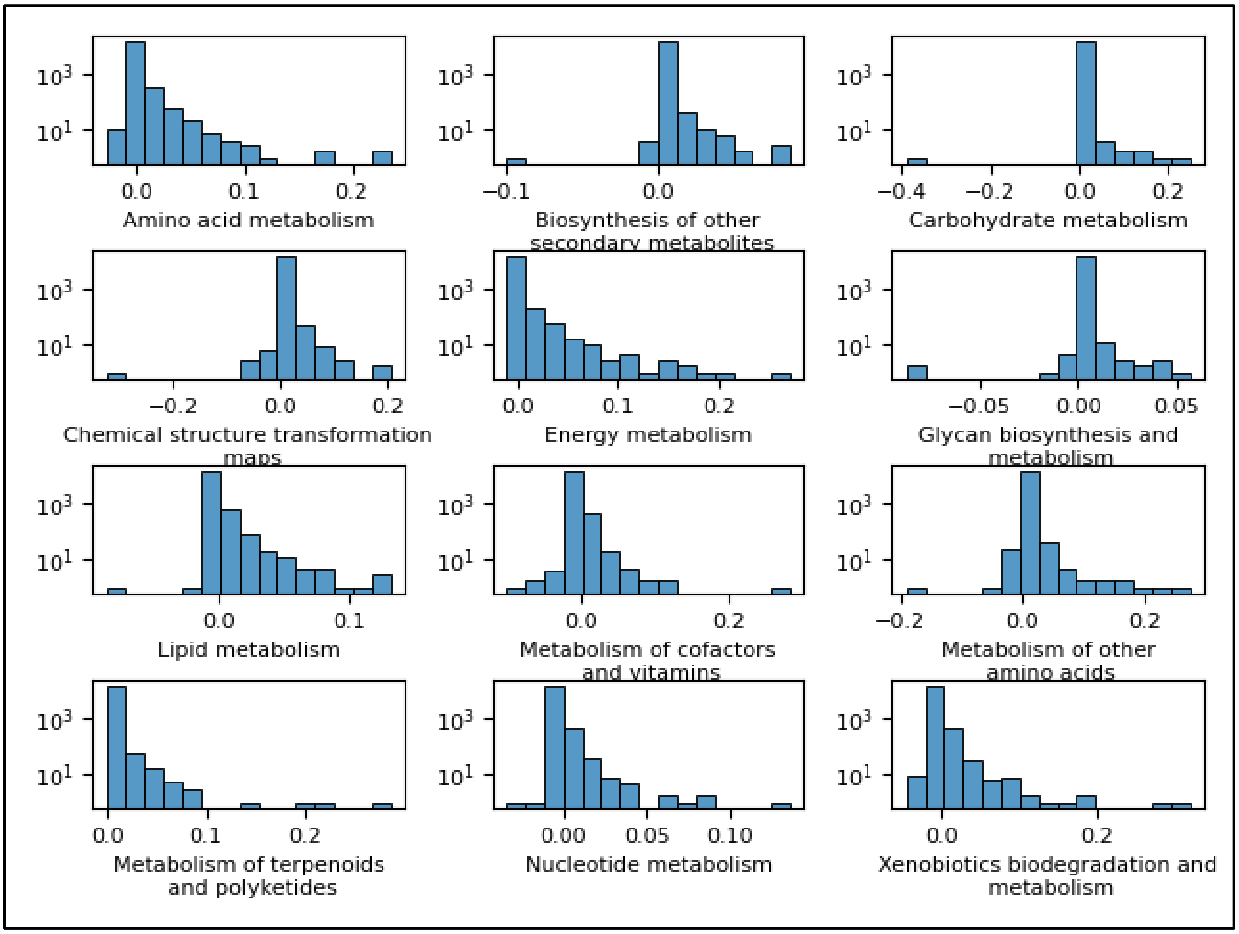


Table S4 – Pathway category proportions in each dataset.

| Dataset | Pathway category | Proportion |
|---|---|---|
| Full | Amino acid metabolism | 0.1075 |
| | Biosynthesis of other secondary metabolites | 0.2615 |

| | | |
|---|---|---|
| | Carbohydrate metabolism | 0.0904 |
| | Chemical structure transformation maps | 0.0769 |
| | Energy metabolism | 0.0304 |
| | Glycan biosynthesis and metabolism | 0.0570 |
| | Lipid metabolism | 0.1191 |
| | Metabolism of cofactors and vitamins | 0.0966 |
| | Metabolism of other amino acids | 0.0482 |
| | Metabolism of terpenoids and polyketides | 0.1923 |
| | Nucleotide metabolism | 0.0297 |
| | Xenobiotics biodegradation and metabolism | 0.1652 |
| Non-ambiguous | Amino acid metabolism | 0.1109 |
| | Biosynthesis of other secondary metabolites | 0.2769 |
| | Carbohydrate metabolism | 0.0876 |
| | Chemical structure transformation maps | 0.0811 |
| | Energy metabolism | 0.0308 |
| | Glycan biosynthesis and metabolism | 0.0388 |
| | Lipid metabolism | 0.1079 |
| | Metabolism of cofactors and vitamins | 0.0976 |
| | Metabolism of other amino acids | 0.0480 |
| | Metabolism of terpenoids and polyketides | 0.1942 |
| | Nucleotide metabolism | 0.0310 |
| | Xenobiotics biodegradation and metabolism | 0.1752 |
| Unfiltered | Amino acid metabolism | 0.1117 |
| | Biosynthesis of other secondary metabolites | 0.2517 |
| | Carbohydrate metabolism | 0.0965 |
| | Chemical structure transformation maps | 0.0725 |
| | Energy metabolism | 0.0397 |
| | Glycan biosynthesis and metabolism | 0.0578 |
| | Lipid metabolism | 0.1265 |
| | Metabolism of cofactors and vitamins | 0.1055 |
| | Metabolism of other amino acids | 0.0567 |
| | Metabolism of terpenoids and polyketides | 0.1851 |
| | Nucleotide metabolism | 0.0317 |
| | Xenobiotics biodegradation and metabolism | 0.1695 |

Table S7 – Valid score counts less than 300.

| **Model** | **Dataset** | **Test Set** | **Pathway Category** | **Metric** | **Valid Score Count** |
|---|---|---|---|---|---|
| Multi-layer Perceptron | Full | Ambiguous | Chemical structure transformation maps | Recall | 229 |
| | | | Nucleotide metabolism | F1 Score | 219 |

| | | | | | |
|---|---|---|---|---|---|
| | | | | Recall | 181 |
| | | | | Precision | 127 |
| | | | Xenobiotics biodegradation and metabolism | F1 Score | 264 |
| | | | | Recall | 215 |
| | | | | Precision | 99 |
| Random Forest | Full | Ambiguous | Biosynthesis of other secondary metabolites | Precision | 159 |
| | | | Chemical structure transformation maps | Recall | 229 |
| | | | Energy metabolism | Precision | 69 |
| | | | Metabolism of other amino acids | Precision | 184 |
| | | | Nucleotide metabolism | F1 Score | 194 |
| | | | | Recall | 181 |
| | | | | Precision | 81 |
| | | | Xenobiotics biodegradation and metabolism | F1 Score | 270 |
| | | | | Recall | 215 |
| | | | | Precision | 142 |
| | Non-ambiguous | Ambiguous | Metabolism of other amino acids | Precision | 200 |
| | | | Nucleotide metabolism | Precision | 1 |
| XGBoost | Full | Ambiguous | Chemical structure transformation maps | Recall | 229 |
| | | | | Precision | 149 |
| | | | Energy metabolism | Precision | 269 |
| | | | Nucleotide metabolism | F1 Score | 190 |
| | | | | Recall | 181 |
| | | | | Precision | 182 |
| | | | Xenobiotics biodegradation and metabolism | Recall | 215 |
| | | | | Precision | 240 |

Figure S2 – MCC By test set for each pathway category for the XGBoost model trained on the full dataset.

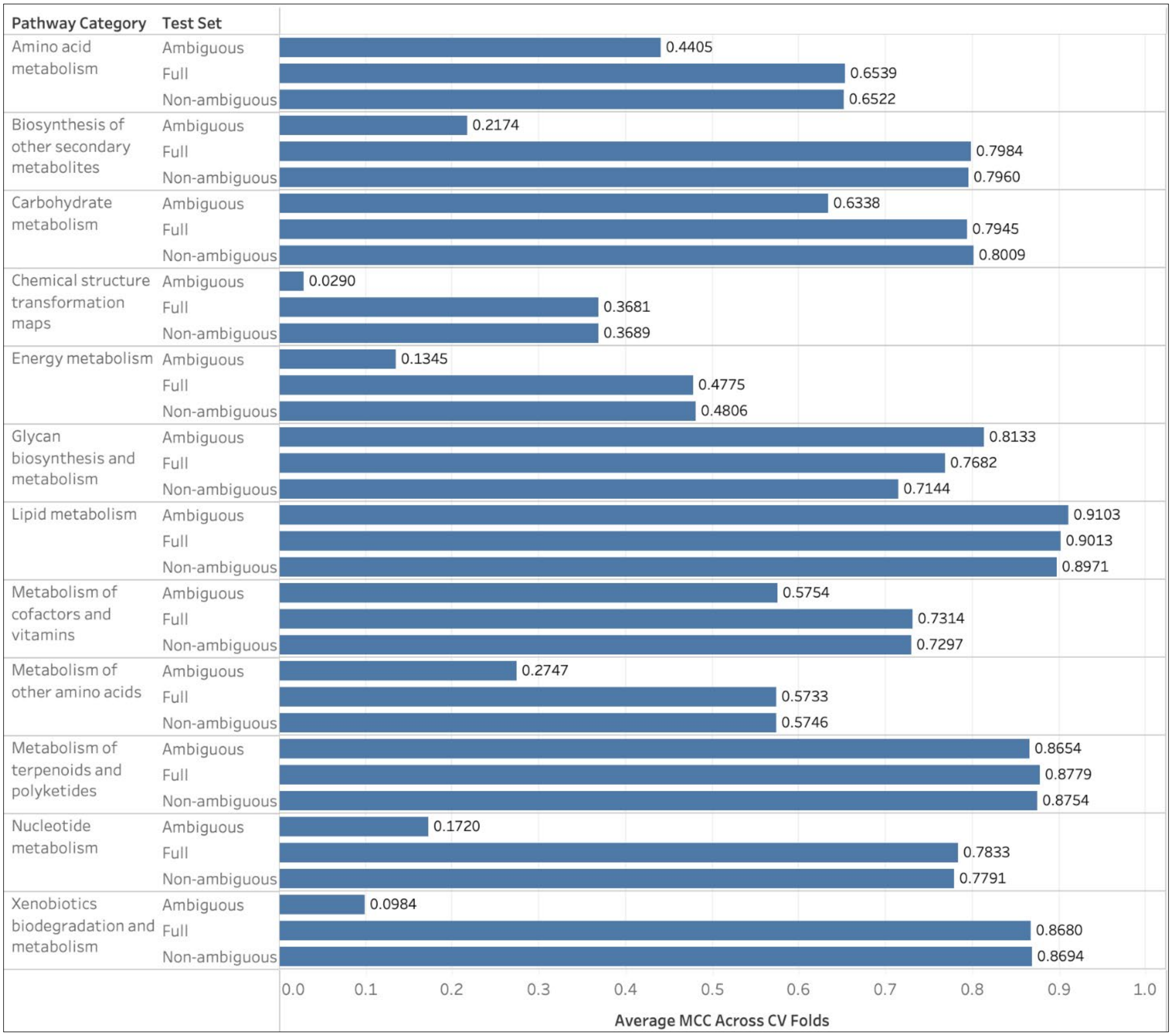


Figure S3 – Upset plot showing overlap between pathway categories of their top 50 most important features.

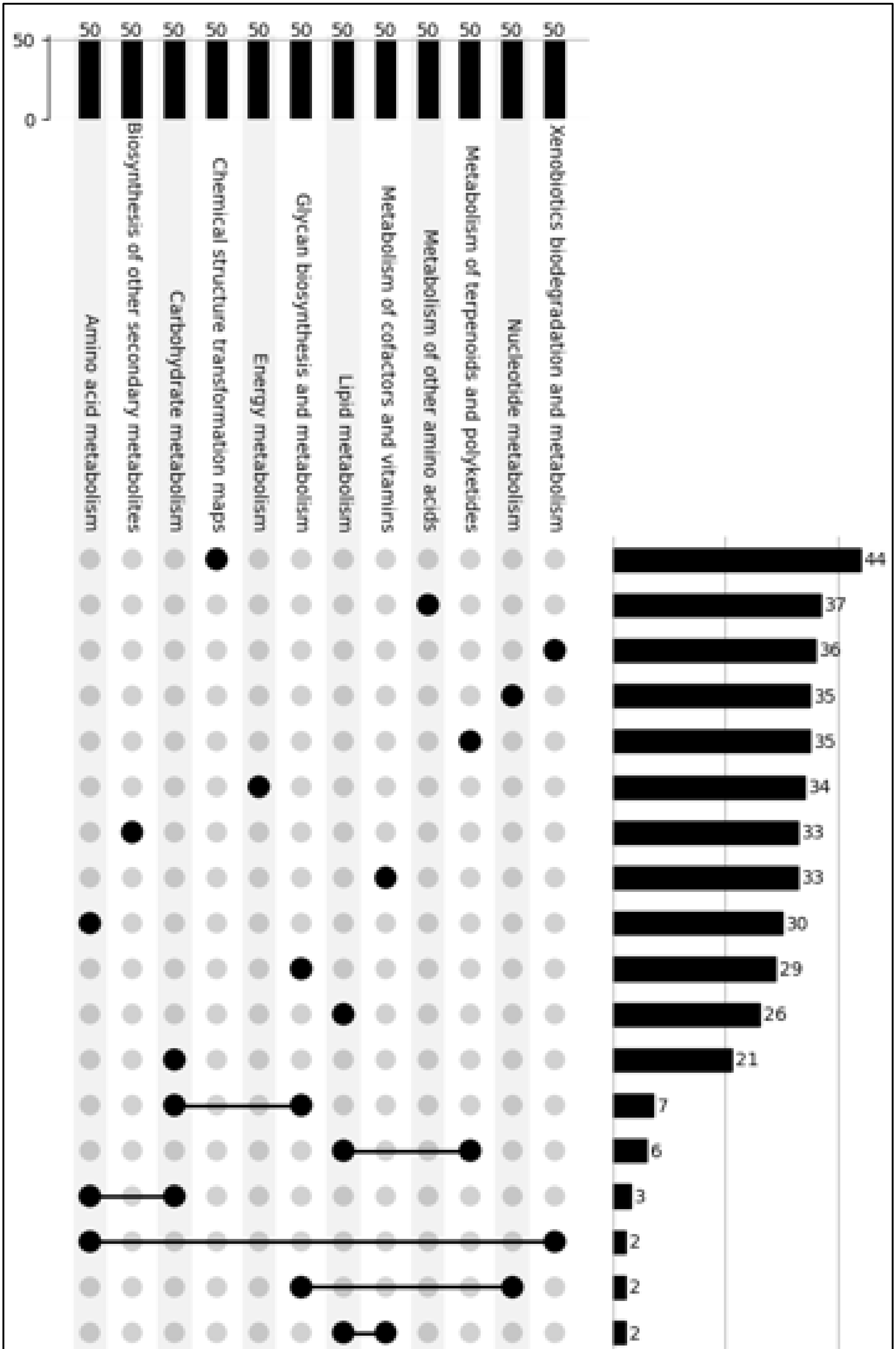
50
0
50
Amino acid metabolism
Biosynthesis of other secondary metabolites
Carbohydrate metabolism
Chemical structure transformation maps
Energy metabolism
Glycan biosynthesis and metabolism
Lipid metabolism
Metabolism of cofactors and vitamins
Metabolism of other amino acids
Metabolism of terpenoids and polyketides
Nucleotide metabolism
Xenobiotics biodegradation and metabolism
44
37
36
35
35
34
33
33
30
29
26
21
7
6
3
2
2
2
